# Antibody Responses to Melanoma Helper Peptide Vaccines May Enhance Antigen Opsonization Through Formation of Immune Complexes and are Modulated by Vaccine Adjuvants

**DOI:** 10.1101/2025.11.24.690236

**Authors:** Emily G. Ashkani, Anna M. Dickinson, Walter C. Olson, Justin J. Taylor, Craig L. Slingluff

## Abstract

**Background:** Vaccines targeting melanoma antigens can elicit CD8^+^ T cell responses, but a growing body of work suggests CD4^+^ T cells also play a role in tumor control. Induction of CD4^+^ cells may also support B cells in producing tumor antigen-specific antibodies (Abs). We investigated Abs induced by vaccination with a cocktail of six class II MHC-restricted melanoma peptides (6MHP) and the effect of adjuvant type on Ab isotypes. We hypothesized that the vaccines would induce Abs that respond to different epitopes on individual peptides and that IgG subclass distribution varies with different vaccine adjuvants.

**Methods:** Sera from patients who received a 6MHP vaccine were evaluated with enzyme-linked immunosorbent assays to map epitopes for polyclonal Ab responses to synthetic melanoma peptides (n=8). IgG subclasses of Ab responses to 6MHP were assessed in patients who received one of 4 adjuvants (Incomplete Freund’s Adjuvant (IFA) alone, IFA + polyICLC, IFA + systemic mCy, or IFA + polyICLC + systemic mCy) to characterize IgG subclass distribution (n=2-14). Comparisons were evaluated using a Mann-Whitney rank sum test.

**Results:** Epitope mapping revealed that at least 50% of patients exhibiting Ab responses to melanoma peptides had responses to two or more epitopes on the same peptide, suggesting polyclonal antibody responses. Serum evaluation for IgG isotypes showed predominant induction of IgG1 and IgG3. Mean total IgG was highest when IFA and polyICLC were used in combination. Patients who received TLR3 agonist polyICLC had significantly higher concentrations of total IgG, IgG1, and IgG3 compared to patients who did not receive polyICLC.

**Conclusions:** These findings suggest that vaccine-induced Abs may respond to multiple epitopes within the same peptide, which may support creation of large antigen-Ab complexes, with promise to facilitate antigen uptake and presentation. Abs were predominantly IgG1 and IgG3, which are optimal for binding complement and supporting Ab-dependent cellular cytotoxicity. The findings also show that adding polyICLC to IFA can significantly enhance Ab responses. Collectively, this work underscores the immunologic potential of peptide-induced Abs and the importance of adjuvant selection in cancer vaccine design.

## 1. BACKGROUND

The ability of cancer vaccines to induce or expand T cell responses to melanoma antigens offers promise for clinical activity as monotherapy or in conjunction with checkpoint blockade therapy and other immune therapies.^1–12^ Most cancer vaccines are designed to induce CD8^+^ cytotoxic T cells specific for tumor antigens, but a growing body of work highlights the ability of CD4^+^ helper T cells to control human cancers.^13–16^ While the induction of tumor antigen-specific CD4^+^ helper T cells could result in direct tumor reactivity, the other main function of CD4^+^ T cells is to support antibody (Ab) production by B cells. Thus, peptide vaccines can also induce tumor antigen-specific Abs,^17,18^ which may further support antitumor responses. However, clinical trials of peptide vaccines have largely ignored the presence and function of induced Abs, or the vaccine strategies that may best support strong antibody responses and antibody class switching.

We have previously reported that vaccination with 6 melanoma helper peptides (6MHP) induced circulating 6MHP-specific IgG responses which were associated with significantly improved patient survival.^19^ The induction of 6MHP-specific Abs could support opsonization and increased antigen uptake and presentation by dendritic cells (DCs), particularly if these Abs are polyclonal and form large immune-complexes (ICs).^20^ It is also important to consider the unexplored potential role for vaccine adjuvants to enhance or inhibit Ab responses to tumor antigens. Presently, there is no consensus about optimal vaccine adjuvants for inducing T cell responses,^21^ and little is known about their effect on Ab responses to peptide antigens. It is known that toll-like receptor (TLR) agonists can have direct and indirect effects on B cell survival and function.^22–28^ Thus, the choice of adjuvants used with vaccination is likely to affect Ab response. A previous study showed that adding a TLR3 agonist to incomplete Freund’s adjuvant (IFA) increased antibody and T cell responses to overlapping long peptides from the cancer-testis antigen NY-ESO-1.^18^ However, how IFA alone or in combination with a TLR3 agonist affects IgG subclass distribution remains unexamined. IgG isotypes vary in complement-binding and antigen-neutralization capabilities;^29^ so, it is important to understand IgG subclass distribution changes with vaccine adjuvants. Abs to the intracellular proteins represented by 6MHP may not have direct antitumor activity through Ab-dependent cellular cytotoxicity (ADCC) or complement-dependent cytotoxicity (CDC). However, direct antitumor activity of Abs to intracellular proteins has been demonstrated in other settings, which may be explained by transient surface expression.^30–32^ IgG1 and IgG3 have the highest affinity for protein antigens and most effectively form immune complexes,^33^ so their predominant induction could result in enhanced vaccine-antigen uptake by DCs.^34,35^ Further, vaccine-induced IgG could bind to the full-length melanocytic differentiation peptides upon their release from lysed melanoma cells, resulting in epitope spreading as these peptides get taken up by DCs and presented to CD8^+^ T cells.^36^ Therefore, identifying Ab-specific epitopes and optimized adjuvants could guide future enhancements of anti-cancer immune responses.^33,37^

In the present study, we evaluated sera from melanoma patients who received the 6MHP vaccine to determine the nature of Abs induced by vaccination with melanoma peptides, and to determine the effects of adjuvant on subclass distribution of IgG. We hypothesized that vaccination with 6MHP vaccine induces Abs that bind two or more different epitopes within individual peptides, which could support the formation of large immune complexes, thus enhancing antigen uptake. We also hypothesized that addition of a TLR agonist to the vaccine emulsion would increase overall IgG induction and induce isotypes with potent complement activation (IgG1 and IgG3).

## 2. MATERIALS AND METHODS

### 2.1 Patient sample preparation

Patient samples were acquired from clinical trials Mel41 (NCT00089219)^2^ and Mel63 (NCT02425306).^38^ In Mel41, patients diagnosed with stage IIIB-IV melanoma were vaccinated weekly (t=6) with 6MHP. 6MHP vaccines contain a cocktail of 6 different peptides of 14-23 amino acids in length, as previously reported (**Table S1**).^2^ Briefly, peptides were administered with 110 μg GM-CSF with 1 mL Montanide ISA-51 adjuvant (IFA, Seppic, Inc., Paris, France/Fairfield, NJ) at either 200 μg (n=12), 400 μg (n=12), or 800 μg (n=13) for each of 6 vaccine administrations. Of the 37 patients vaccinated, 8 patients with previously identified favorable Ab responses were selected for testing against the overlapping peptide sequences (**Table S2**).

In trial Mel63, patients were treated in an adaptive design, with enrollment to one of 4 arms (A-D), where 6MHP vaccines included IFA (A), IFA with polyinosinic-polycytidylic acid stabilized with polylysine and carboxymethylcellulose (polyICLC, Hiltonol, Oncovir, Washington, D.C.) (B), IFA with systemic metronomic cyclophosphamide (mCy) (C), or all three in combination as previously reported (D) **(Table 1)**.^38^ Briefly, all patients were vaccinated with 200 μg of the 6MHP pool **(Table S1)** in either IFA (Montanide ISA-51VG, Seppic Inc.) alone or IFA with polyICLC, with or without oral mCy **(Table 1)**. Vaccines were administered on days 1, 8, 15, 36, 57, and 78, and blood was collected at several time points. Of the 48 patients enrolled in the study, 26 patients were selected for evaluation of isotype/subtype specificity **(Table S3)**.

**Table 1.**
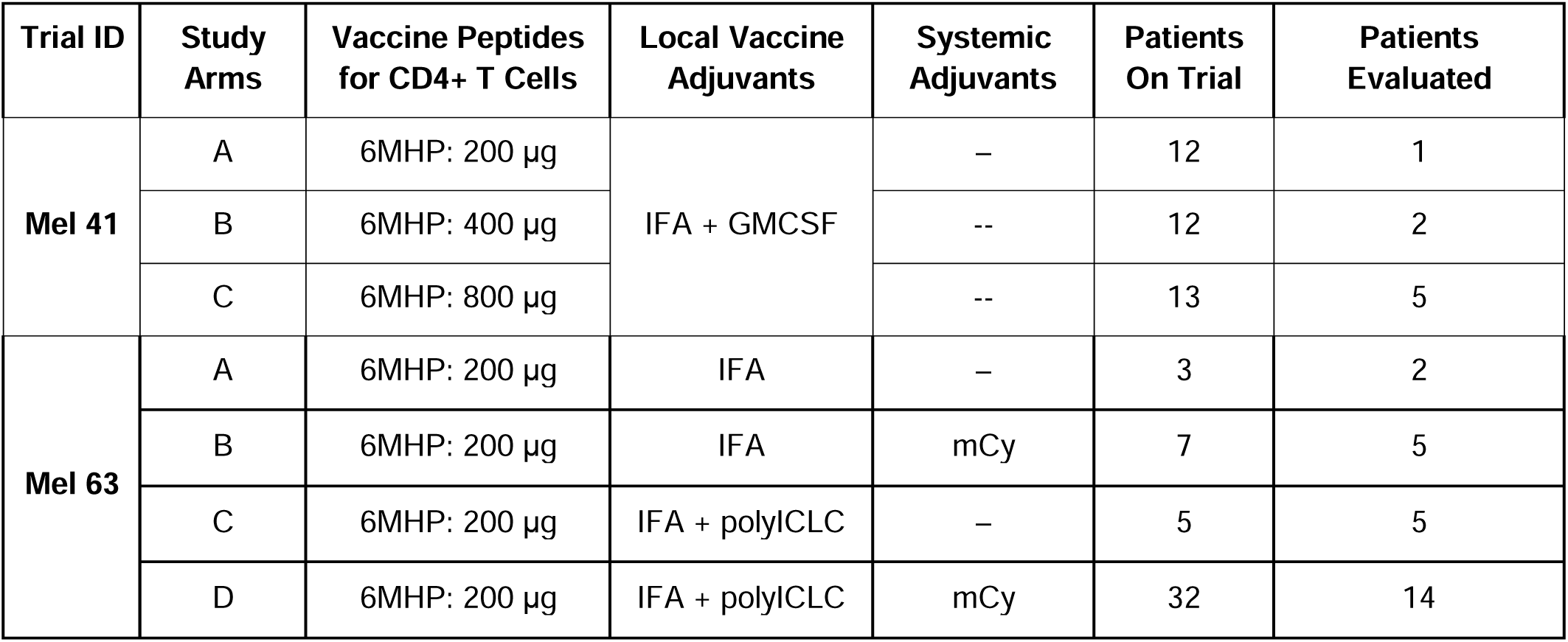
Clinical Trial Information for Mel41 and Mel63.

### 2.2 Peptides

The peptides in the 6MHP pool were synthesized in GMP conditions for the vaccine trials themselves. Peptides representing overlapping sequences within the full-length peptides were synthesized by GenScript (Piscataway, New Jersey, USA). Of the six peptides in the 6MHP vaccine, the highest Ab response rates were observed for the three longest peptides (gp100_44-59_, Tyrosinase_386-406,_ Melan-1/MART-1_51-73_)^19^; thus, these were selected for this study **(Table 2)**. Overlapping peptides were synthesized, representing eleven amino acids in length and spanning each of the peptides with five overlapping residues.

**Table 2.**
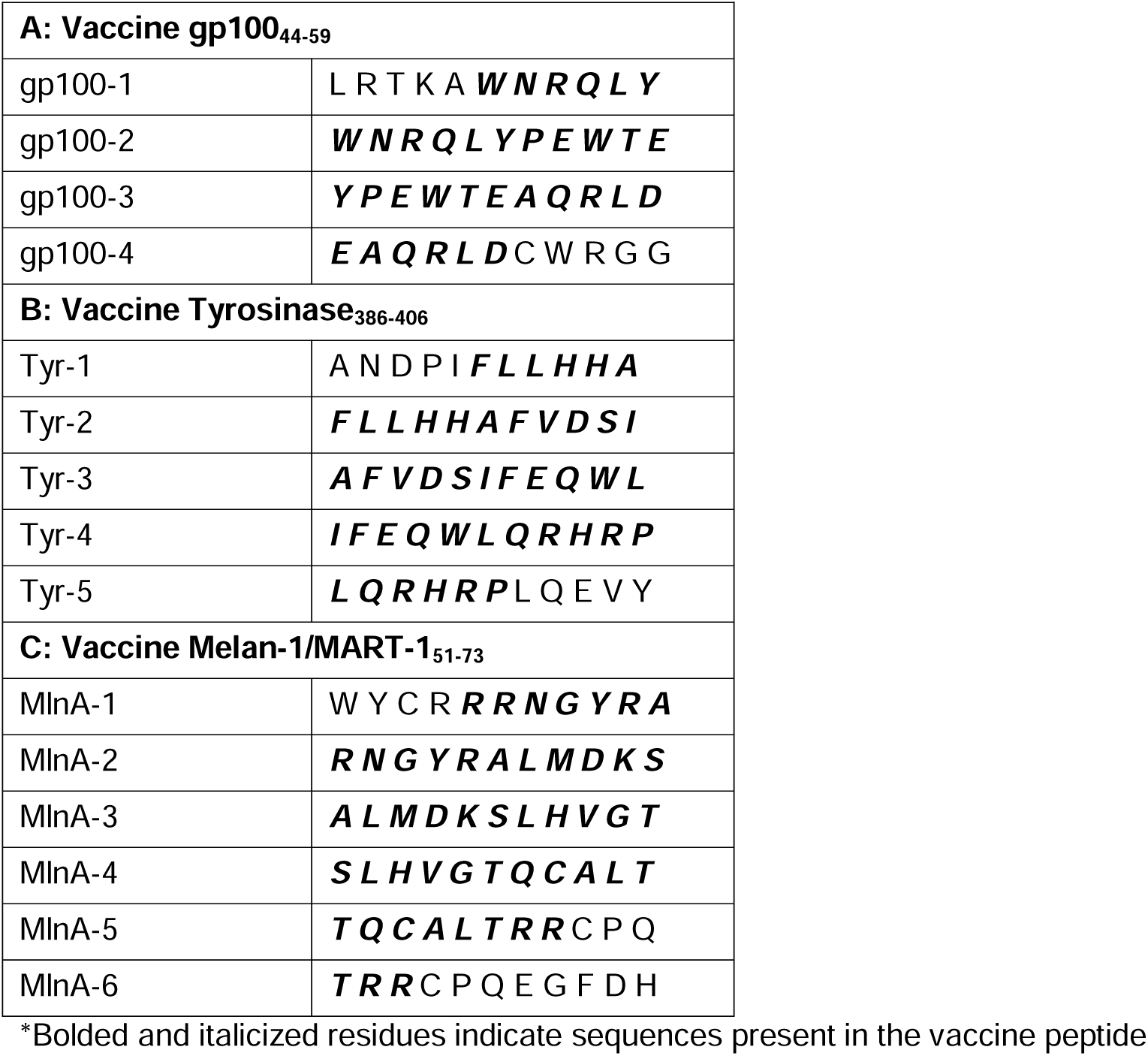
List of overlapping peptides and their sequences*.

### 2.3 Detecting responses to overlapping peptides by ELISA

Direct enzyme-linked immunosorbent assays (ELISAs) were used to evaluate patient sera from the Mel41 trial (week 12 post-vaccination) for total IgG responses to overlapping 11- mer peptides using described methods.^19,39^ Briefly, 96-well half-area polystyrene plates (Corning, Cat. No. 3690) were coated with peptide preparations: individual full-length peptides as a positive control (**Table S1**), 11-mer overlapping peptide sequences (**Table 2**), or HIV-GAG polymerase derived 20-amino acid peptide (GAG_293-312_, FRDYVDRFYKTLRAEQASQE) as a negative control. All peptide concentrations were adjusted to 1.67 μg/mL in a bicarbonate buffer (pH 9.4; Sigma-Aldrich, Cat No. C3041) and 30 μL were added to each well. The plates incubated overnight at 4 °C and were then washed four times with wash buffer (phosphate-buffered saline (PBS)/0.1% Tween 20). The plates were blocked with blocking buffer containing 5% non-fat (NF) dry milk and 0.1% sodium azide (pH 7.5) in PBS for 1 hour at room temperature. Patient serum was diluted 1:400 in assay buffer (2% normal goat serum (Vector Labs, Cat. No. S-1000, Newark, CA) in blocking buffer) and 30 μL were added to each well. Sera from two normal donors was used as a negative control. The plates were covered and allowed to incubate overnight in a humidified chamber at 4°C. Following incubation, plates were washed several times in wash buffer. 30 uL of alkaline phosphatase (AP) conjugated secondary antibody (goat anti-human IgG, Cat No. 2040-04, Southern Biotechnology Associates) was diluted 1:2000 in assay buffer and 30 uL were added to each well. Plates were allowed to incubate for 1 hour at room temperature. Then, plates were washed three times with wash buffer followed by three times in PBS before 30 μL of AttoPhos AP Fluorescent substrate (Promega, Cat. No. S1000) was added and incubated for 30 minutes at room temperature, light-protected. 15 μL of 3N NaOH was added to each of the wells to stop the reaction.

### 2.4 Measuring IgG subclass distribution by ELISA

Direct ELISAs were used to evaluate patient sera from the Mel63 trial (peak titer, weeks 12, 18, or 26 post-vaccination)^38^ using the described methods.^19,39^ Briefly, 96-well half-area cluster plates (Corning, Cat. No. 3696) were coated with 6MHP (**Table S1**) or the HIV-GAG polymerase-derived 20-amino acid peptide (GAG_293_) as a negative control. All peptide concentrations were adjusted to 1.67 μg/mL in a bicarbonate buffer (pH 9.4; Sigma-Aldrich) and 30 μL were added to each well. The plates were incubated overnight at 4 °C, then washed four times with wash buffer. The plates were blocked with blocking buffer for 2 hours at room temperature. A serial four-fold dilution series of patient serum was prepared in assay buffer, starting with 1:100, and 30 μL of each dilution were then added to the plates in singlets. Sera from two normal donors were used as a negative control, and serum from a patient with known positive responses was used as a positive control. After overnight incubation at 4 °C and subsequent washes with wash buffer, AP-conjugated secondary antibodies (goat anti-human IgG, mouse anti-human IgG1, Cat. No. 9054-04, mouse anti-human IgG2, Cat. No. 9070-04, mouse anti-human IgG3, Cat. No. 9210-04, mouse anti-human IgG4, Cat. No. 9200-04, Southern Biotechnology Associates) were prepared at 1:1000 (IgG1 and IgG3), 1:500 (IgG2 and IgG4) and 1:2000 (IgG total) in assay buffer, and 30 μL of the secondary antibody solutions were added to their assigned wells. The plates were incubated for 1 hour at room temperature. Plates were washed as described above before 30 μL of AttoPhos AP Fluorescent substrate (Promega) was added and incubated for 30 minutes at room temperature, protected from light. 15 μL of 3N NaOH was added to each of the wells to stop the reaction.

### 2.5 Analysis of antibody responses

The SPECTRAmax Gemini EM Fluorescent plate reader (Molecular Devices, Sunnyvale, CA) was used to read the plates at an excitation length of 450/50 nm and emission length of 580/50 nm. The cutoff for positive responses to overlapping peptides was determined to be 2x greater than the background fluorescence level. Background fluorescence was determined by averaging the fluorescence levels of the wells that were coated with the GAG_293-312_ peptide. The background was calculated individually for each plate and then averaged across the three plates.

Positive IgG total and isotype responses to full-length 6MHP peptide were determined to be fluorescence intensities greater than 10x the fluorescence readout of the normal donor serum. A standard curve of fluorescent intensity and concentrations of IgG total and IgG1-4 was generated for each isotype. All plates were corrected for background fluorescence before analysis. Background fluorescence was calculated for each plate by averaging the fluorescence readout of wells that received antigen with no secondary antibody, no antigen with secondary antibody, or normal donor serum. Upper and lower limits of fluorescence intensity were determined based on the minimum and maximum values included in the polynomial curve with a correlation coefficient greater than 0.999. 6MHP-specific IgG total and IgG1-4 concentrations were extrapolated according to the polynomial expression derived from this curve.

### 2.6 Statistical analysis

Comparison of total IgG and subclass concentrations between trial arms was evaluated using a Mann-Whitney rank sum test. All statistical tests were performed using MedCalc Statistical Software version 23.2.7 (MedCalc Software Ltd, Ostend, Belgium; https://www.medcalc.org; 2025).

## 3. RESULTS

### 3.1 Mapping epitope location on peptides

To map the peptide epitopes recognized by serum Abs to Tyr_386_, MelanA_51_, and gp100_44_, sera from patients on the Mel41 trial were assessed using an ELISA. Of the 37 patients on that trial, eight with favorable antibody responses were chosen as a sample population to be assessed for Ab responses in sera at week 12 post-vaccination to the overlapping peptides **(Table 2, Table S2)**. All patients exhibited Ab responses to 6MHP (**Figure 1A-C**). Although there was reactivity to multiple overlapping peptides, the magnitude of reactivity differed among the peptides, as represented by fluorescence levels (**Figure 1A-C**). Of these patients, all eight had positive IgG responses to at least one peptide fragment from tyrosinase_386-406_, while five had positive IgG responses to at least one peptide fragment from Melan-A/MART-1_51-73_, and three had positive responses to at least one peptide fragment from gp100_44-59_ (**Figure 1D-F**). Ab epitopes in Tyrosinase appeared in all fragments except Tyr-1 (**Figure 1D**), while reactivity was exhibited to four of the six segments in MelanA, excluding MlnA-4 and MlnA-5 (**Figure 1E**).

**Figure 1.**
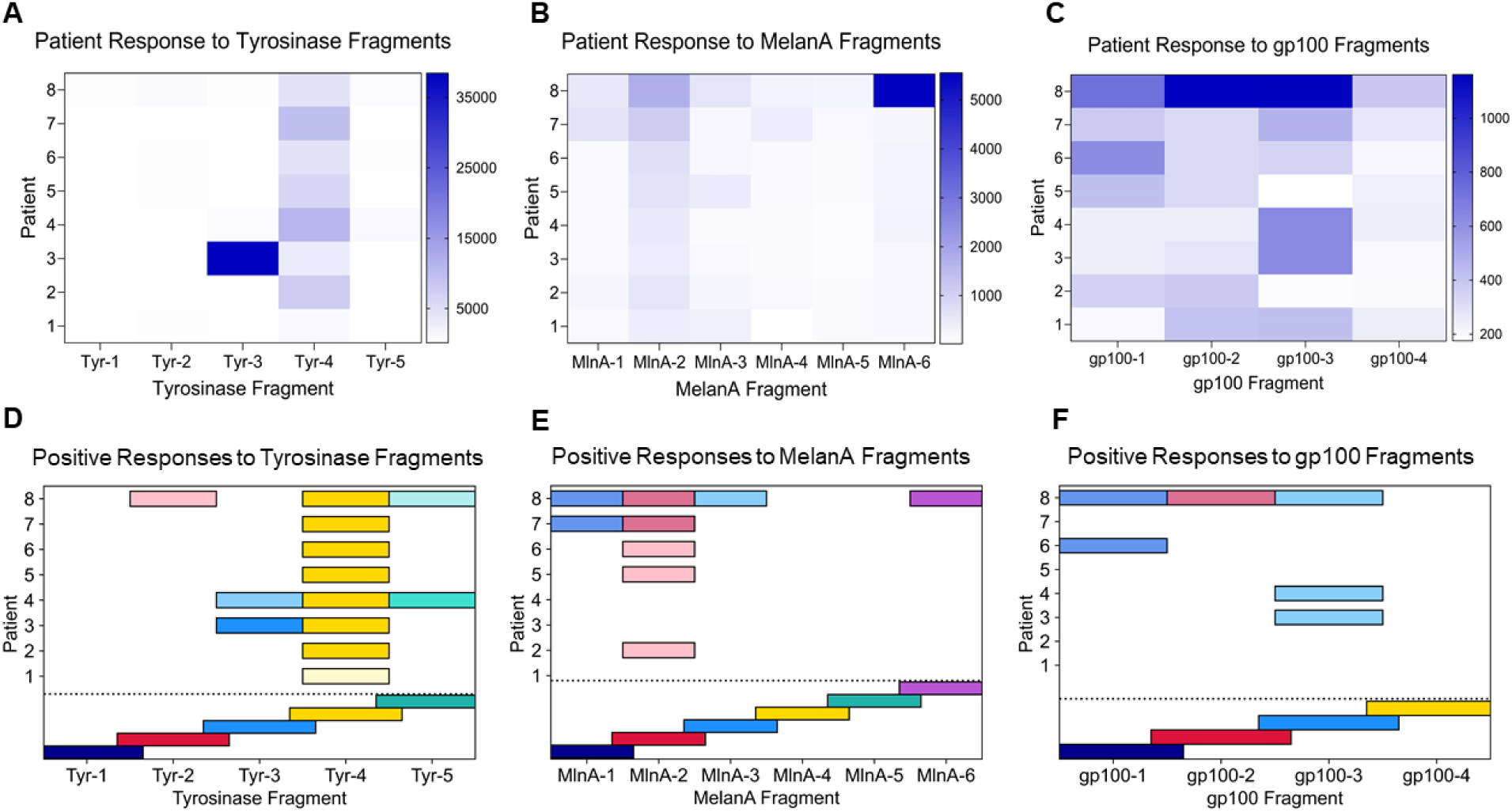
Patient antibody responses to fragments of peptides from 6MHP vaccine. A-C: Heatmaps representing varying degrees of reactivity (in fluorescence units (FU)) to peptide fragments (**A:** Tyrosinase_386-406_ [Tyr], **B:** MelanA_51-73_ [MlnA] for each patient, **C:** gp100_44-59_ [gp100]). Colors reflect the gradient scale on the right side of each heatmap, representing FU (Tyr: 5000-35000, MelanA: 1000-5000, gp100: 200-1000). **D-F:** Positive responses to the overlapping peptides varied among patients, as indicated by the varying intensity gradients. Reference layers at the bottom represent the individual overlapping peptides tested. Data is based on fluorescence units (FU). The lowest fluorescence values considered positive are those greater than two times the average background fluorescence. Colors are scaled to be reflective of ≥2x the average background (lightest), ≥4x the average background, and ≥10x the average background (darkest) for each fragment.

Reactivity to gp100 was observed in three of four segments, excluding gp100-4 (**Figure 1F**). Positive responses to Tyr-4, MlnA-2, and gp100-3 were the most frequent across patients (**Figure 1D-F**). Five patients had IgG responses to at least two separate peptide fragments (patients 3, 4, 7, 8), and two had IgG responses to non-overlapping peptide fragments: patients 4 (Tyr) and 8 (Tyr, MlnA, and gp100) **(Figure 1D-F)**.

### 3.2 Peptide isotype distribution with adjuvant use

To determine the effect of adjuvant on IgG isotype distribution induced by vaccination with helper peptides, IgG subclasses IgG1-IgG4 and total IgG specific to 6MHP were assessed in sera from 26 patients in the Mel63 trial at the time point previously identified as peak titer^38^ (week 12, 18, or 26 post-vaccination) (**Table S3**). Positive total IgG responses were observed in 24 patients (92%) (**Figure 2A**). The predominant 6MHP-specific IgG subclasses were IgG1 (in 50% of responding patients) and IgG3 (in 96% of responding patients) regardless of trial arm (**Figure 2A**). No patients exhibited positive 6MHP-specific IgG2 or IgG4 responses (**Table S3**).

**Figure 2:**
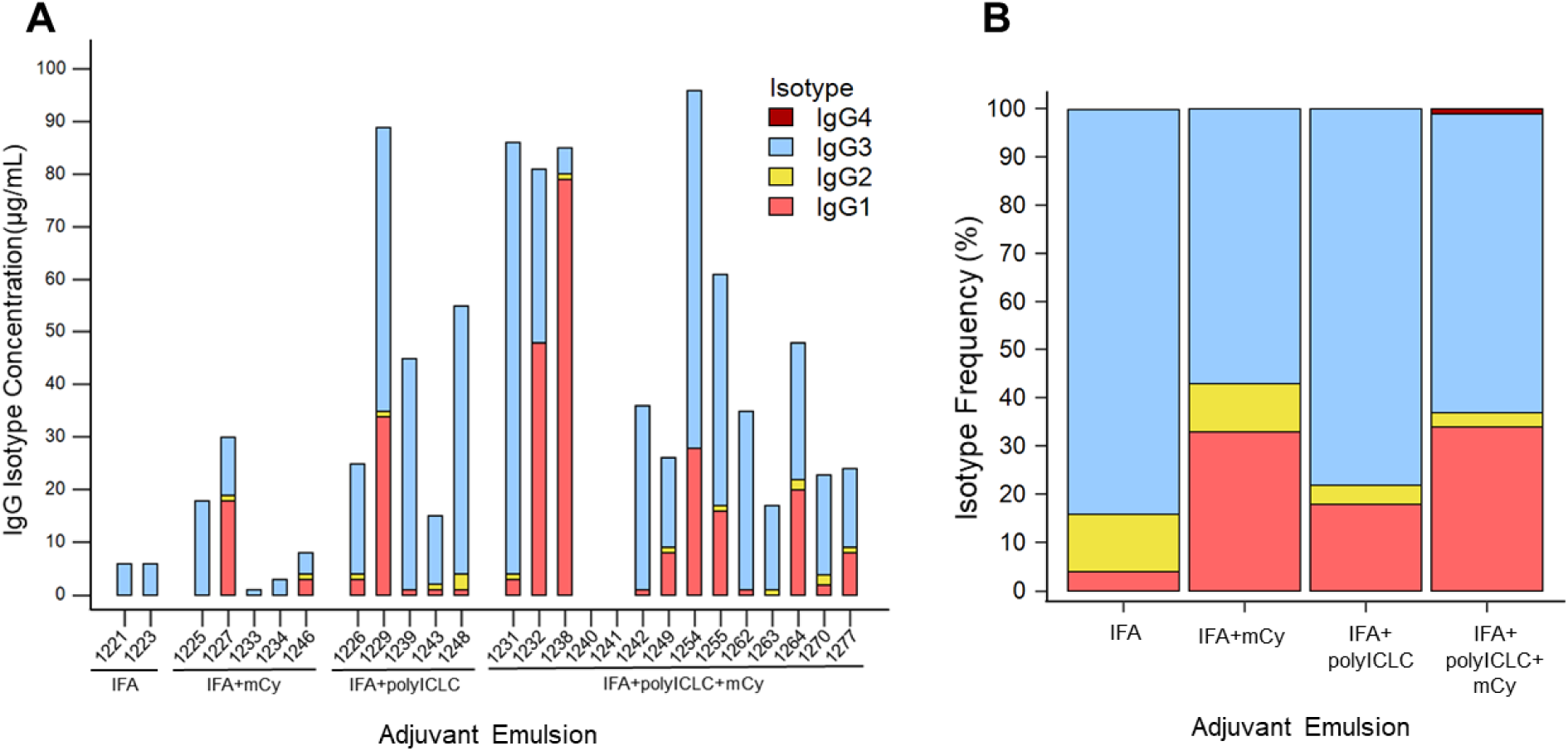
IgG isotype frequencies for each patient and clinical trial arm. Frequencies represent the proportion of each isotype contributing to the sum of all IgG isotypes present: IgG1 (pink), IgG2 (yellow), IgG3 (light blue), and IgG4 (red). **A:** IgG subclass distribution for each patient, separated by trial arm. **B:** IgG subclass distribution for each trial arm. IgG1-4 concentrations were averaged for all patients within each arm before calculating frequency for each trial arm.

Upon separating patients by trial arm, adjuvants appeared to have a modulatory effect on IgG subclasses present in post-vaccination serum. In each trial arm, 6MHP-specific IgG3 was the most abundant isotype detected, averaging from 57% - 84% of the total 6MHP-specific IgG (**Figure 2B**). 6MHP-specific IgG1 was the second most abundant isotype for all arms except for the group that received IFA alone (**Figure 2B**). Instead, 6MHP-specific IgG2 was the second most frequent isotype for patients in the IFA alone arm (**Figure 2B**). In contrast, 6MHP- specific IgG4 was only detectable for patients in the IFA + polyICLC + mCy group and accounted for 1% of IgG across the whole group (**Figure 2B**).

6MHP-specific IgG responses were higher for patients receiving IFA and polyICLC (Arms C and D) than for those receiving 6MHP in IFA alone (**Figure 3A**). 6MHP-specific IgG concentrations were significantly higher for patients receiving IFA + polyICLC (Arm C) compared to patients receiving IFA + mCy (Arm B) (p = 0.016) (Figure 3A). 6MHP-specific IgG was also significantly higher for patients receiving IFA + polyICLC + mCy (Arm D) than for patients receiving IFA alone (Arm A) and IFA + mCy (Arm B) (p = 0.022 and p = 0.0006, respectively) (**Figure 3A**). 6MHP-specific IgG1 was significantly higher in patients receiving IF + polyICLC + mCy (Arm D) compared to patients receiving IFA alone (Arm A) (p = 0.022) (**Figure 3B**). 6MHP-specific IgG3 was significantly elevated for patients receiving IFA + polyICLC, with and without mCy (Arms C and D) compared to patients receiving IFA with mCy (p = 0.016 and p = 0.0061, respectively) (**Figure 3B**).

**Figure 3.**
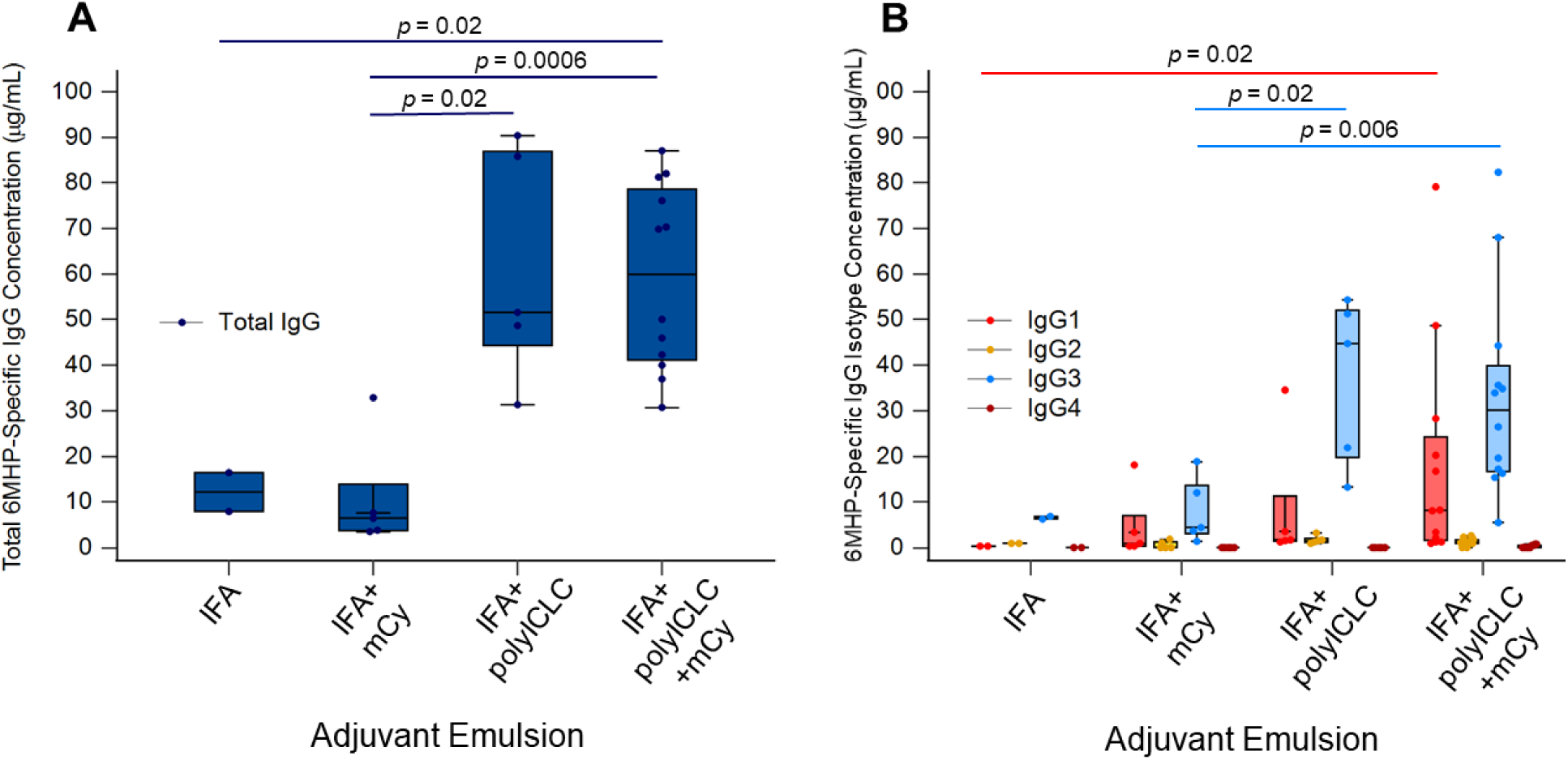
Serum IgG isotypes and total IgG specific for 6MHP vaccines across clinical trial arms. A: Total 6MHP-specific IgG concentration for each trial arm. **B:** 6MHP-specific IgG isotype concentrations (IgG1 (red), IgG2 (yellow), IgG3 (light blue), and IgG4 (maroon)) for each trial arm. Each point represents an individual sample; the center line is the median, the limits are the interquartile range (IQR), and the whiskers extend to the lowest and highest values no greater than 1.5x IQR, with outliers represented by points beyond the whiskers. Significant differences are marked with p values as determined by a Mann-Whitney rank sum test.

Comparing those with and without polyICLC, concentrations of 6MHP-specific IgG, IgG1, and IgG3 were all significantly higher for patients who received polyICLC (Arms C+D vs Arms A+B: p = <0.0001, 0.01, 0.0004, respectively) (**Figure 4A**). Patients who received systemic mCy (Arms B+D) had no significant differences in 6MHP-specific IgG, IgG1, or IgG3 versus those who did not (Arms A+C) (**Figure 4B**). However, IgG1 concentration trended higher for patients receiving mCy (**Figure 4B**).

**Figure 4.**
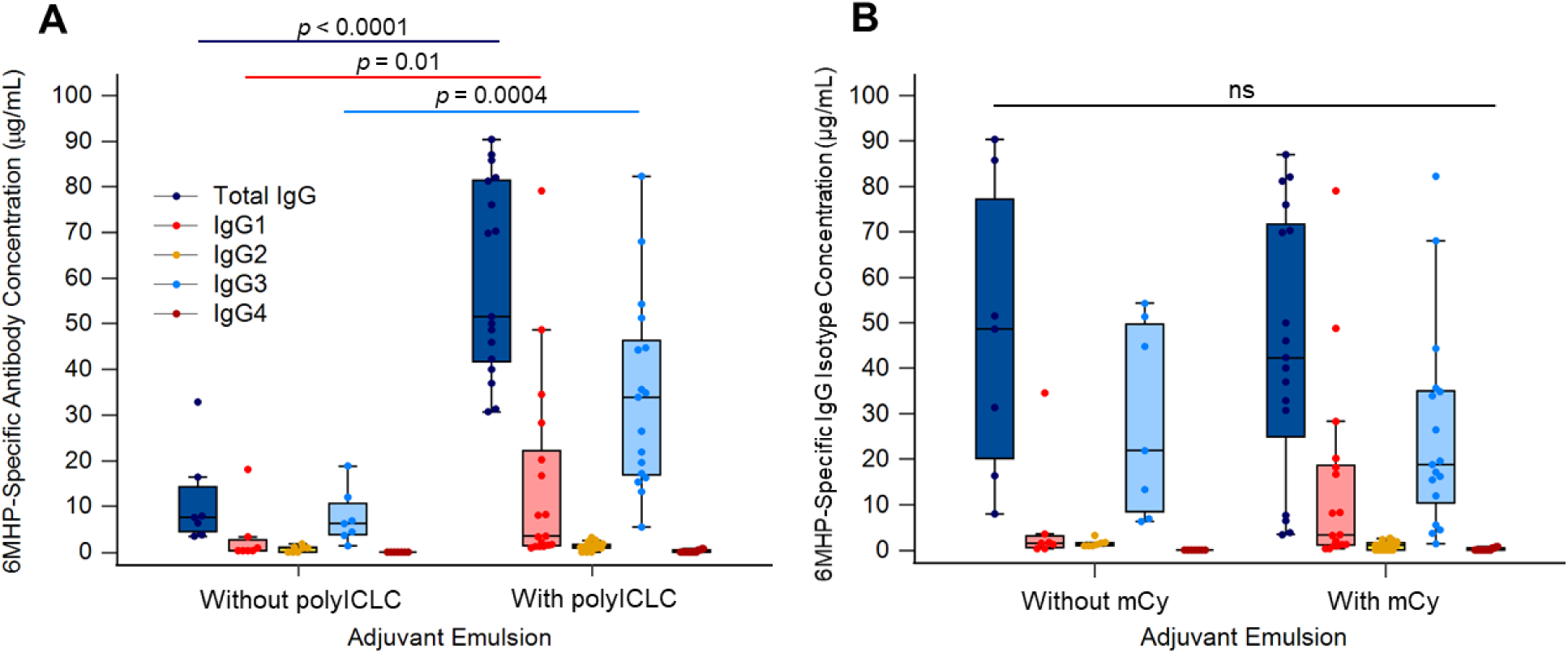
Serum IgG isotypes and total IgG specific for 6MHP vaccines with different adjuvant emulsions. Isotype (IgG1 (pink), IgG2 (yellow), IgG3 (light blue), and IgG4 (red)) concentrations for **A:** patients who received polyICLC (Arms C+D) or did not receive polyICLC (Arms A+B) and **B:** for those who received systemic mCy (Arms B+D) or did not receive mCy: (Arms A+C). Each point represents an individual sample; the center line is the median, the limits are the interquartile range (IQR), and the whiskers extend to the lowest and highest values no greater than 1.5x IQR, with outliers represented by points beyond the whiskers. Significant differences are marked with p values as determined by a Mann-Whitney rank sum test; ns: not significant.

## 4. DISCUSSION

We have found that vaccination with 6MHP in an emulsion with IFA induces antibodies to those peptides in over 90% of patients. IgG1 and IgG3 were detected as the predominant IgG subclasses and are associated with higher affinity for protein antigens and a greater ability to form immune complexes.^33^ Most patients had antibodies specific to more than one epitope on individual peptides, but only a quarter of the patients exhibited reactivity to non-overlapping fragments (**Figure 1D-F**). This means there is a possibility of forming large immune complexes by binding to multiple sites along peptides, which could facilitate uptake and presentation by antigen presenting cells (APCs).^40^ However, this may only occur for a subset of patients. We have also found that adding a TLR3 agonist to the vaccine adjuvant enhances total IgG Ab production as well as enhances both IgG1 and IgG3, supporting our hypothesis (**Figure 4A**).

In our prior studies, we found that of the proteins incorporated into the pool of 6MHP, the HLA-DR-restricted peptides derived from melanocytic differentiation proteins (Tyrosinase, gp100, MelanA) induced Ab responses at the highest frequencies.^19^ Here, we used synthetic peptide segments containing 11 amino acids **(Table 2)** from the original 16-23 amino acid sequences **(Table S1)** to map the regions containing epitopes to which the induced antibodies were responding. These peptide segments overlapped by 5 amino acids, allowing for the detection of epitopes spanning segments. For the overlapping peptides from Tyrosinase, binding sites appeared in all segments except for Tyr-1 **(Figure 1D)**. MelanA also exhibited reactivity in 4 out of the 6 segments, except for MlnA-4 and MlnA-5 (**Figure 1E**). In gp100, reactivity appeared in 3 out of 4 segments, excluding gp100-4 (**Figure 1F**). Five of the eight patients had positive responses to two or more epitopes. It is possible that reactivity to adjacent epitopes is indicative of Ab response to the same epitope rather than a polyclonal response since each peptide segment overlaps by five amino acids. However, reactivity was observed in non-overlapping residue sequences for two patients **(Figure 1D-F)**. Therefore, we can conclude that two or more distinct epitopes are recognized by Ab in at least some patient sera, but that this is not common for most patients. A polyclonal response would be important as it could allow for the formation of large ICs, targeting DC Fc-gamma receptors to introduce these antigens into the cross-presentation pathway.^40^ This is known to induce strong CD8^+^ T cell responses, but can also allow for CD8^-^ DCs to effectively cross-present exogenous antigens.^40^ Thus, polyclonal 6MHP-specific antibody responses may enhance antigen uptake and thereby increase CD8^+^ T cell immune response to the 6MHP vaccine.^40^ Vaccine-induced B cell and DC responses could also lead to epitope spreading to other melanoma antigens, as the Abs could bind to full-length peptides after they are released from dead melanoma cells.^36^ This would allow for uptake and presentation of new melanocytic differentiation protein peptides by DCs, inducing new populations of melanoma antigen-specific CD8^+^ T cells. Epitope spreading is important for successful immunotherapy, and activated B cells are the main APC responsible for expanding T cell responses.^36^ Therefore, the induction of melanoma antigen-specific B cells and DCs may ultimately augment tumor immunity. Future studies will include a finer specificity assessment of the epitope sequences using shorter length peptides and peptides with alanine substituted for individual residues.

The combination of adjuvants in conjunction with TLR agonists in the 6MHP vaccine was found to be correlated with IgG response. The patients selected for serological evaluation received vaccinations with IFA alone, IFA with polyICLC, IFA with systemic mCy, or all three in combination. Patient serum samples were selected from the time point yielding the highest titer of 6MHP-specific antibodies based on previously completed analysis^38^ (12, 18, or 26 weeks) (**Table S3**). Each serum specimen was tested for total IgG and subclasses IgG1-IgG4 to the 6MHP peptides. Positive IgG responses were observed in 24 of the 26 patients evaluated (92% response rate) (**Figure 2A**). Of those who responded, IgG1 and IgG3 were the predominant subclasses represented (in 50% and 96% of responding patients, respectively). IgG2 and IgG4 responses were negative for all patients (**Table S3**). In healthy donors, the most abundant IgG isotype is typically IgG1, making up approximately 65% of circulating IgG, followed by IgG2 at 26%.^41^ Here, IgG3 was the most frequently detected subclass across study arms (**Figure 2B**). We observed IgG1 frequencies of 4-34% across trial arms, and IgG3 frequencies of 57-84% across trial arms (**Figure 2B**). The quantification of IgG in this study was restricted to only IgG specific for the 6MHP vaccine. As IgG3 is strongly immune activating,^33^ we expect it to be the most abundant. Prior studies have also demonstrated IgG3’s role in immune response against proteins or polypeptide antigens, supporting their presence in response to the 6MHP peptides.^37^ Additionally, it has been found that IL-21, primarily produced by CD4^+^ T cells, plays a role in IgG1 and IgG3 production by inducing class switch recombination events in naïve IgG**^-^** B cells.^42^ When considered alongside our findings of predominant IgG1 and IgG3 induction, this suggests that vaccine-induced CD4^+^ T cells can augment B cells to form a robust adaptive immune response against the vaccine antigens.

Adjuvant emulsion was shown to be associated with changes in concentrations of total IgG, IgG1, and IgG3 (**Figure 3A-B**). IFA alone has previously been shown to increase CD8^+^ T cell immune response rate,^43^ and effectively induced 6MHP-specific antibodies in the present study.

The largest increases in IgG were observed in patients who received TLR3 agonist polyICLC in addition to IFA (**Figure 4A**). Significant changes were not observed when comparing patients who received mCy versus those who did not (**Figure 4B**). Future studies will focus on characterizing the effects of vaccine adjuvants on isotype class switching, as well as the longevity of IgG subclass responses induced by the 6MHP peptide vaccine to further understand the impact of adjuvants on antibody-mediated immune responses.

## 5. CONCLUSION

We have shown that Abs produced in response to the 6MHP vaccine are specific for at least two epitopes on each tested helper peptide, which may support the development of large antigen-Ab complexes that could enhance antigen presentation by DCs. Induction of large antigen-Ab complexes may be especially beneficial in the case of short peptides that present on MHC I, as the DC cross-presentation pathway allows for these exogenous antigens to be recognized by CD8^+^ T cells. This may also allow for epitope spreading to induce new 6MHP- specific CD8^+^ T cell populations, if these Abs bind to full-length protein released from lysed melanoma cells and facilitate DC uptake and presentation of new epitopes. We also show that polyICLC used in addition to IFA can significantly enhance Ab responses to the 6MHP vaccine. Collectively, this work highlights the immunologic potential of peptide-induced Abs and the importance of adjuvant selection in cancer vaccine design.

## DECLARATIONS

### Ethics Approval and Consent to Participate

The clinical trials Mel41 and Mel63 were performed with IRB (#10464 and 17860, respectively), and FDA (BB-IND #10825) approval and were registered with Clinicaltrials.gov (NCT00089219 and NCT02425306, respectively).

## Consent for Publication

All authors have revised the manuscript and consented to its publication.

## Availability of data and material

All data relevant to the study are included in the article or uploaded as supplementary information.

## Authors’ contributions

EGA: data analysis, figure creation, manuscript writing

AMD: data collection, data analysis, manuscript writing

WCO: data collection, data analysis, experimental planning

JJT: data analysis, manuscript revision

CLS: conceptual design, experimental planning, manuscript revision, funding acquisition

### List of Abbreviations

IFA: Incomplete Freund’s Adjuvant
polyICLC: polyinosinic-polycytidylic acid stabilized with poly-lysine and carboxymethylcellulose
mCy: Metronomic cyclophosphamide
6MHP: 6 melanoma helper peptides
ELISA: enzyme-linked immunosorbent assay
Ab: antibody
IgG: immunoglobulin G
IC: immune complex
TLR: toll-like receptor
GMP: good manufacturing practices
GM-CSF: granulocyte macrophage colony stimulating factor
NF: non-fat
PBS: phosphate buffered saline
AP: alkaline phosphatase
MHC: major histocompatibility complex
DC: dendritic cell
APC: antigen presenting cell
ADCC: antibody dependent cellular cytotoxicity

