## Supplementary material for "Antibody Responses to Melanoma Helper Peptide Vaccines May Enhance Antigen Opsonization Through Formation of Immune Complexes and are Modulated by Vaccine Adjuvants": Table S1

### Table S1. Peptide sequences from 6MHP Vaccine

| **Allele** | **Sequence** | **Epitope** |
|---|---|---|
| HLA-DR4 | AQNILLSNAPLGPQFP | $Tyrosinase_{56-70}$ |
| HLA-DR15 | FLLHHAFVDSIFEQWLQRHRP | $Tyrosinase_{386-406}$ |
| HLA-DR4 | RNGYRALMDKSLHVGTQCALTRR | Melan-A/MART-$1_{51-73}$ |
| HLA-DR11 | TSYVKVLHHMVKISG | MAGE-$3_{281-295}$ |
| HLA-DR13 | LLKYRAREPVTKAE | MAGE-1,2,3,$6_{121-134}$ |
| HLA-DR1 & HLA-DR4 | WNRQLYPEWTEAQRLD | $gp100_{44-59}$ |
