## Supplementary material for "Antibody Responses to Melanoma Helper Peptide Vaccines May Enhance Antigen Opsonization Through Formation of Immune Complexes and are Modulated by Vaccine Adjuvants": Table S2

**Table S2. Mel41 Serum Samples Included in Study**

| Patient # | VMM | Trial Arm | Serum used (weeks post-vaccination) |
|---|---|---|---|
| 8 | VMM 871 | B | Week 12 |
| 7 | VMM 729 | C | Week 12 |
| 6 | VMM 719 | C | Week 12 |
| 5 | VMM 701 | A | Week 12 |
| 4 | VMM 699 | B | Week 12 |
| 3 | VMM 683 | C | Week 12 |
| 2 | VMM 625 | C | Week 12 |
| 1 | VMM 485 | C | Week 12 |
