## Supplementary material for "Antibody Responses to Melanoma Helper Peptide Vaccines May Enhance Antigen Opsonization Through Formation of Immune Complexes and are Modulated by Vaccine Adjuvants": Table S3

**Table S3. Mel63 Patients Included in Study and IgG Concentrations (μg/mL)**

| Patient | Trial Arm | Serum used (weeks post-vaccination) | IgG1 | IgG2 | IgG3 | IgG4 | IgG Total |
|---|---|---|---|---|---|---|---|
| VMM1221 | A | Week 18 | 0.32 | 0.96 | 6.8 | 0 | 16 |
| VMM1223 | A | Week 18 | 0.28 | 0.96 | 6.3 | 0 | 8.0 |
| VMM1225 | B | Week 18 | 0.31 | 0 | 19 | 0 | 3.4 |
| VMM1227 | B | Week 18 | 18 | 1.8 | 12 | 0 | 33 |
| VMM1233 | B | Week 18 | 0.28 | 0 | 1.4 | 0 | 3.8 |
| VMM1234 | B | Week 18 | 0.89 | 0 | 3.6 | 0 | 6.5 |
| VMM1246 | B | Week 18 | 3.3 | 1.1 | 4.5 | 0 | 7.6 |
| VMM1226 | C | Week 26 | 3.5 | 1.6 | 22 | 0 | 49 |
| VMM1229 | C | Week 26 | 35 | 1.5 | 54 | 0 | 90† |
| VMM1239 | C | Week 26 | 1.5 | 0.86 | 45 | 0 | 86 |
| VMM1243 | C | Week 18 | 1.7 | 1.2 | 13 | 0 | 31 |
| VMM1248 | C | Week 18 | 1.2 | 3.2 | 51 | 0 | 52 |
| VMM1231 | D | Week 26 | 3.3 | 1.1 | 82 | 0 | 87 |
| VMM1232 | D | Week 26 | 49 | 0 | 34 | 0 | 70 |
| VMM1238 | D | Week 26 | 79 | 1.2 | 5.5 | 0 | 76 |
| VMM1240 | D | Week 26 | 0.93 | 0 | 0 | 0 | 0.10 |
| VMM1241 | D | Week 18 | 1.8 | 0 | 0 | 0 | 0.05 |
| VMM1242 | D | Week 18 | 1.2 | 0 | 36 | 0 | 50 |
| VMM1249 | D | Week 26 | 8.0 | 1.5 | 17 | 0 | 30 |
| VMM1254 | D | Week 26 | 28 | 0 | 68 | 0.46 | 81 |
| VMM1255 | D | Week 18 | 17 | 1.1 | 44 | 0 | 82 |
| VMM1262 | D | Week 18 | 1.2 | 0.86 | 34 | 0 | 37 |
| VMM1263 | D | Week 12 | 0.98 | 1.1 | 16 | 0.48 | 42 |
| VMM1264 | D | Week 18 | 20 | 2.3 | 26 | 0.70 | 70 |
| VMM1270 | D | Week 12 | 2.1 | 2.5 | 20 | 0.77 | 40 |
| VMM1277 | D | Week 12 | 8.3 | 1.8 | 15 | 0.47 | 46 |
| Number (%) positive by arm | A | | 0 (0%) | 0 (0%) | 2 (100%) | 0 (0%) | 2 (100%) |
| | B | | 2 (40%) | 0 (0%) | 4 (80%) | 0 (0%) | 5 (100%) |
| | C | | 2 (40%) | 0 (0%) | 5 (100%) | 0 (0%) | 5 (100%) |
| | D | | 8 (57%) | 0 (0%) | 12 (86%) | 0 (0%) | 12 (86%) |
| Overall | | | 12 (46%) | 0 (0%) | 23 (88%) | 0 (0%) | 24 (92%) |

*Shaded boxes are ones that did not meet the threshold of >10x the FU of normal donor serum to be considered a positive response

*Concentrations of 0 were below the detectable range

†IgG total was not properly detected for this patient, so the sum of IgG subclasses is used in place of total IgG value
